# Potential For Applying Continuous Directed Evolution To Plant Enzymes

**DOI:** 10.1101/2020.08.26.265678

**Authors:** Jorge D. García-García, Jaya Joshi, Jenelle A. Patterson, Lidimarie Trujillo-Rodriguez, Christopher R. Reisch, Alex A. Javanpour, Chang C. Liu, Andrew D. Hanson

**Affiliations:** Horticultural Sciences Department, University of Florida, Gainesville, FL, USA; Department of Microbiology and Cell Science, University of Florida, Gainesville, FL, USA; Department of Biomedical Engineering, University of California, Irvine, Irvine, CA, USA; Department of Chemistry, University of California, Irvine, Irvine, CA, USA; Department of Molecular Biology and Biochemistry, University of California, Irvine, Irvine, CA, USA

**Author notes:** These authors contributed equally to this work (Jorge D. García-García, Jaya Joshi).

**Keywords:** Protein engineering, Synthetic biology, Linear plasmids, Error-prone polymerases, CRISPR/Cas9, Directed evolution

## Abstract

Plant evolution has produced enzymes that may not be optimal for maximizing yield and quality in today’s agricultural environments and plant biotechnology applications. By improving enzyme performance, it should be possible to alleviate constraints on yield and quality currently imposed by kinetic properties or enzyme instability. Enzymes can be optimized faster than naturally possible by applying directed evolution, which entails mutating a target gene *in vitro* and screening or selecting the mutated gene products for the desired characteristics. Continuous directed evolution is a more efficient and scalable version that accomplishes the mutagenesis and selection steps simultaneously *in vivo* via error-prone replication of the target gene and coupling of the host cell’s growth rate to the target gene’s function. However, published continuous systems require custom plasmid assembly, and convenient multipurpose platforms are not available. We discuss two systems suitable for continuous directed evolution of enzymes, OrthoRep in *Saccharomyces cerevisiae* and EvolvR in *Escherichia coli*, and our pilot efforts to adapt each system for high-throughput plant enzyme engineering. To test our modified systems, we used the thiamin synthesis enzyme THI4, previously identified as a prime candidate for improvement. Our adapted OrthoRep system shows promise for efficient plant enzyme engineering.

## INTRODUCTION

Enzyme evolution is naturally driven by beneficial mutations that are selected over many generations, but this inherently slow and random process does not necessarily produce optimally adapted enzymes, especially for new environments or applications [1,2]. In fact, many enzymes – in plants as in other organisms – are merely ‘good enough’ to support metabolism. For example, enzymes with sub-par kinetic properties can limit key metabolic pathways, and the frequent replacement of short-lived enzymes can divert resources from biomass synthesis and other productive uses [2,3]. There is thus much potential for plant improvement by reducing enzyme inefficiencies [4,5], a goal that can be advanced by applying approaches from outside plant biology.

Synthetic biology (SynBio) offers a suite of such approaches. SynBio lies at the intersection between discovery-driven biology and goal-oriented engineering and brings a fresh perspective to plant improvement by adding to the breeding toolbox [6,7]. SynBio has enormous potential in enzyme engineering, ranging from improving existing catalytic activities to creating new ones [7–9]. Early enzyme engineering in the 1980s focused on rational (re)design by site-specific mutagenesis [10]. However, evolving enzymes through semi-rational mutagenesis depends on knowledge of structure-function relationships that is often unavailable [11] and is time- and labor-intensive, particularly when multiple mutations are needed to achieve the desired improvement. A classic example of the challenges of enzyme improvement by site-specific mutation is Rubisco, ribulose-1,5-bisphosphate carboxylase/oxygenase [12]. Attempts to improve Rubisco’s carboxylase activity in plants have been complicated by the number of possible target residues, with roles ranging from conformational flexibility [13] to protein-protein interactions [14,15] to effector/inhibitor binding [16,17]. It is therefore close to impossible to predict an optimal set of mutations to maximize catalytic efficiency.

### Directed Evolution

An alternative to rational design approaches is directed evolution [18], which takes a cue from nature by harnessing the power of random mutagenesis. Directed evolution typically begins by diversifying a target gene by introducing random mutations to generate large libraries of mutated sequences. The genes encoded in these mutated sequences are then expressed, and the enzymes screened *in vitro* or *in vivo* for the desired properties [19]. The advantage of directed evolution is that its outcomes are not constrained by knowledge about the target enzyme. Rather, it is an ‘act first, ask questions later’ approach in which mutations are made iteratively or combinatorially in the target gene until the desired outcome is obtained.

The effects of cumulative mutations on enzyme activity can be visualized as a ‘fitness landscape’ in which compounding beneficial changes lead to fitness peaks and detrimental changes cause fitness valleys [20] (Figure 1). Given the existence of fitness valleys, reaching an optimal peak may require ‘valley crossing’ through intermediate steps that are deleterious [21,22]. Traversing valleys can occur by relaxing selective pressure or by introducing multiple mutations simultaneously, effectively skipping over the intermediate, less-fit steps. Directed evolution can cope with fitness valleys by introducing multiple mutations during each round of mutagenesis, but its separate screening step is labor-intensive, which limits throughput capacity. Tuning selection pressure is likewise labor-intensive.

**Figure 1.**
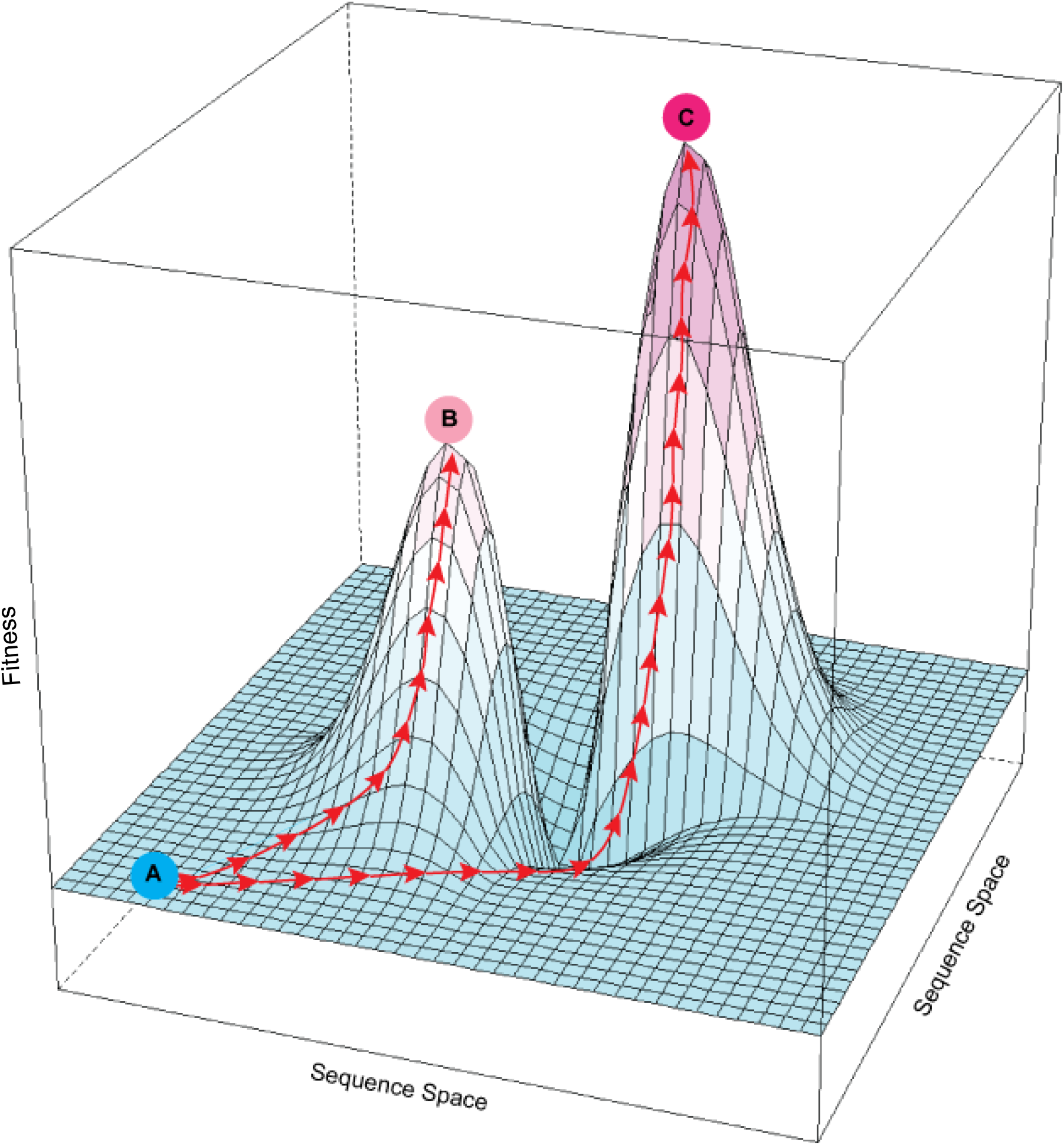
Rugged fitness landscape with multiple fitness peaks. The *x* and *y* axes represent the multidimensional sequence space defined by different possible sites and identities of mutations in an enzyme; the *z*-axis is fitness as a function of sequence. Red arrows show possible evolutionary paths created by mutation and selection. An enzyme at suboptimal fitness level A could potentially reach local fitness peak B or global optimal peak C. Because the path from A to C passes through a fitness valley it is more difficult to follow under selection than the continuously upward path from A to B.

### Continuous Directed Evolution

Continuous directed evolution is a ‘next-generation’ version of directed evolution that accomplishes the mutagenesis and screening steps simultaneously by continuously applying selection pressure throughout the mutation process [23–25]. In continuous directed evolution, the function of the target gene is typically coupled to the growth rate of the host microbe by forcing the microbe to rely on the target gene, e.g. by complementing the loss of an essential host activity with a plasmid-borne target gene. Using this method, improved target genes are immediately identifiable with no need for additional screening [26–28]. Also, tracking enzyme function *in vivo* avoids possible artifacts of *in vitro* screening assays, which may be poor facsimiles of physiological conditions. A particular advantage of continuous directed evolution is its scalability, which allows exploration of many trajectories through a fitness landscape *simultaneously* (which is challenging with classical directed evolution, see above). Some continuous directed evolution platforms are also compatible with automation, which furthers scalability and industrialization [29].

The first successful continuous directed evolution system for proteins was Phage-Assisted Continuous Evolution (PACE), in which M13 bacteriophage carry a plasmid-borne target gene that is iteratively propagated and mutated within infected *E. coli* host cells [23]. The target gene’s function is coupled to the induction of an essential bacteriophage virulence factor, pIII, such that more active target gene variants result in increased phage infectivity and propagation. Although novel pIII induction mechanisms are being developed to adapt PACE for broader applications [25], the utility of PACE for metabolic enzyme evolution remains limited, there being no generalizable way to link pIII induction to enzyme traits such as kinetic characteristics or half-life.

Fortunately, continuous directed evolution systems suitable for metabolic enzyme improvement have now been introduced. Below, we outline the *Saccharomyces cerevisiae* OrthoRep and *Escherichia coli* EvolvR systems; each has unique and innovative features that avoid some of the limitations of PACE.

### OrthoRep

The yeast OrthoRep system was developed from the p1 and p2 linear cytoplasmic plasmids of *Kluyveromyces lactis* [26,30–32]. The p1-specific DNA polymerase TP-DNAP1 was transferred from p1 to a nuclear plasmid, leaving a backbone into which a target gene can be introduced (Figure 2A); p2 remains in its native state, encoding its own DNA polymerase (TP-DNAP2) and the machinery needed to transcribe both p1 and p2 [32]. For continuous directed evolution, error-prone versions of TP-DNAP1 have been engineered; these polymerases introduce mutations into the p1-borne target gene without increasing the genomic mutation rate (Figure 2A). Error-prone TP-DNAP1s have mutation rates as high as 1 × 10^−5^ substitutions per base, which is up to 10^5^ times that of the host genome [26,32]. The ~10^5^-fold mutational acceleration, combined with serial passaging, enables rapid, continuous evolution of the target gene entirely *in vivo.* OrthoRep is compatible with high-throughput, automated systems, including some that can impose diverse and adjustable selection pressures to help cross fitness valleys [29].

**Figure 2.**
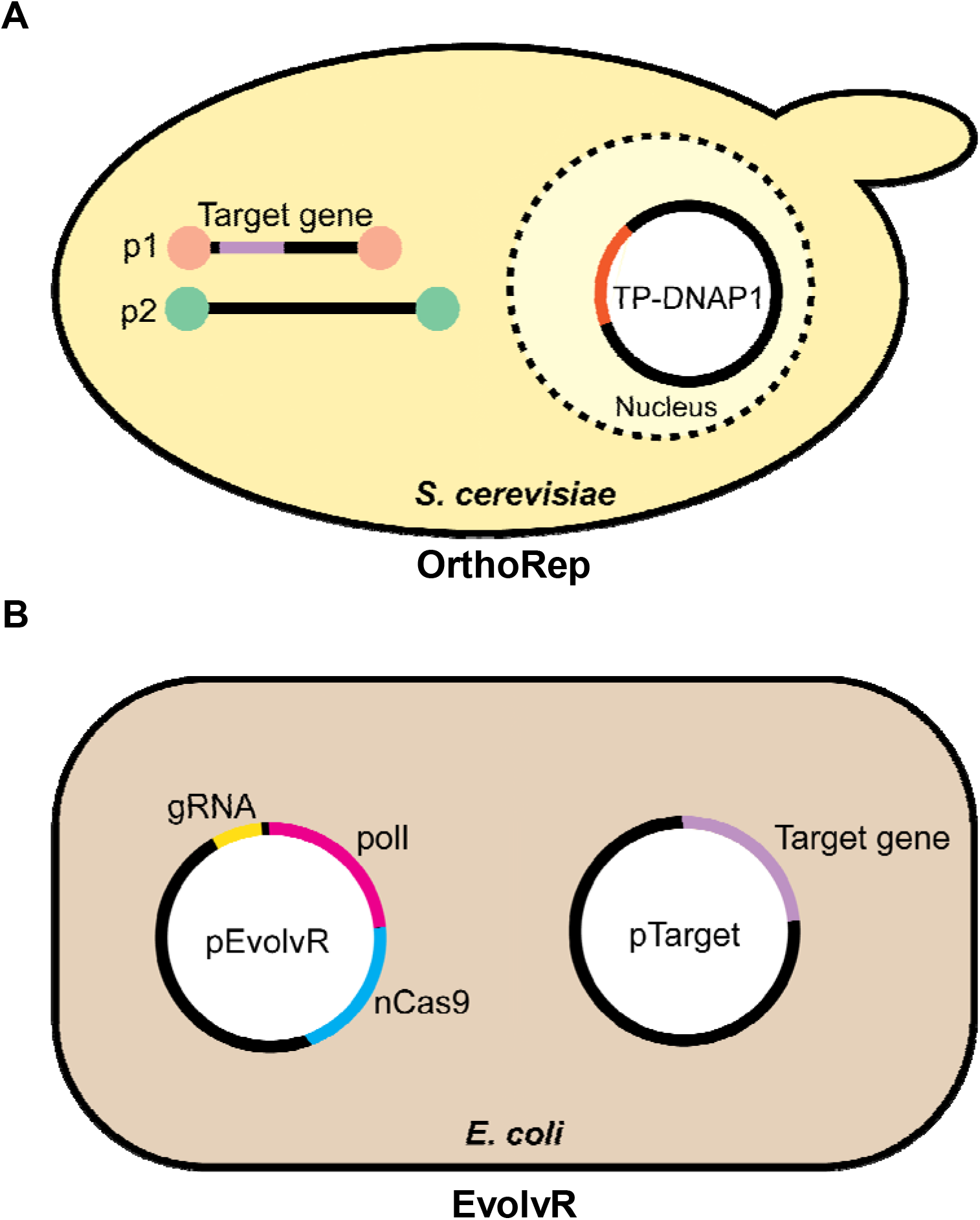
The OrthoRep and EvolvR continuous directed evolution systems. Schematic overviews of the two continuous evolution systems evaluated in this study. **(A)** The yeast OrthoRep system consists of an error-prone DNA polymerase, encoded by a nuclear plasmid, that replicates cytoplasmic linear plasmid p1, and only p1. The p1 plasmid is engineered to express the target gene. Cytoplasmic linear plasmid p2 carries the machinery needed to transcribe the genes on both p1 and p2, and a high-fidelity DNA polymerase that replicates p2 alone. **(B)** The *E. coli* EvolvR system comprises the pEvolvR plasmid, encoding an nCas9 nickase-error-prone DNA polymerase I fusion plus a guide RNA, and pTarget encoding the target gene.

### EvolvR

The *E. coli* EvolvR system uses a fidelity-reduced variant of *E. coli* DNA polymerase I (PolI3M or PolI5M) fused to a CRISPR-guided nickase, nCas9 (derived from *Streptococcus pyogenes* Cas9), which directs the polymerase to a specific location on the target gene [27] (Figure 2B). EvolvR is a two-plasmid system in which one plasmid (pEvolvR) harbors the PolI–nCas9 fusion plus a single-guide RNA (sgRNA) and the other plasmid (pTarget) carries the target gene. The EvolvR system offers a targeted mutation rate of ~10^−5^ to 10^−6^ mutations per nucleotide per generation averaged over a 56-bp window, with the highest rates closest to the nick site [27].

### Limitations of Continuous Directed Evolution Systems

The application of OrthoRep, EvolvR, or other continuous directed evolution systems to high-throughput improvement of enzymes is subject to certain (mostly obvious) constraints, as follows. (i) As noted above, the target enzyme typically needs to be essential to the growth of the platform cells so that the growth rate depends on the enzyme’s activity. (ii) The target enzyme must be expressed mainly in soluble form to avoid selecting for mutations that merely increase solubility in the alien platform cell milieu instead of improving the desired characteristic immediately. (iii) To maximize applicability, the system should function in minimal or defined growth media. While OrthoRep was developed using defined media, EvolvR was developed using rich medium and its compatibility with minimal media was not explored.

To date, OrthoRep and EvolvR have only been used to evolve microbial enzymes, and for just two types of applications (antibiotic resistance and thermal adaptation) [27,29,32]. Furthermore, particularly for OrthoRep, the target gene plasmid construct is tailor-made and cannot be easily repurposed for different genes of interest.

The aim of this work was two-fold: (i) to adapt OrthoRep to facilitate simple, high-throughput cloning of target genes; and (ii) to establish the potential of OrthoRep and EvolvR to improve the characteristics of metabolic enzymes, which – for EvolvR – required use of minimal medium. We chose the thiamin thiazole synthesis enzyme THI4 as the target for these proof-of-concept studies. Plant and yeast THI4s are suicide enzymes that self-inactivate after a single catalytic cycle because they obtain the sulfur atom needed to form the thiazole product by destroying an active-site cysteine residue. Such THI4s are consequently energetically expensive to operate [3,33–35]. In contrast, certain prokaryotic THI4s are truly catalytic, i.e. perform multiple reaction cycles; these enzymes use sulfide as sulfur source [36,37]. *Thermovibrio ammonificans* THI4 (TaTHI4) belongs to this group [35,38]. However, like other characterized prokaryotic THI4s, TaTHI4 prefers anaerobic, high-sulfide conditions, making it ill-suited for function in plant cells. Our long-term goal is therefore to evolve the non-suicidal TaTHI4 to function efficiently in plant-like conditions (aerobic, low sulfide). To explore the feasibility of this goal we adapted OrthoRep and EvolvR systems to complement thiamin auxotrophic yeast or *E. coli* strains with native yeast THI4 or TaTHI4, respectively.

## MATERIALS AND METHODS

### Chemicals, enzymes, and oligonucleotides

Chemicals were purchased from Sigma–Aldrich (St. Louis, MO) unless otherwise stated. Phusion® High-Fidelity DNA polymerase and restriction enzymes (New England Biolabs, Ipswich, MA) were used for PCR reactions and gene cloning. Oligonucleotide primers were obtained from Eurofins Genomics LLC (Louisville, KY) and are listed in Supplementary Table S1. All constructs were sequence-verified.

### OrthoRep

#### Construction of multipurpose p1 integration vector

In previous work, target gene expression in yeast was enhanced by placing a poly-A tail/self-cleaving ribozyme (RZ) sequence downstream of the target gene [39]. We therefore built a multipurpose p1 integration vector from 172-YTK-P4 (containing the poly-A/RZ self-cleaving ribozyme cassette) and GR306 (used as backbone, with the fluorescent reporter *mKate2* in its cloning region), as shown in Figure 3. We excised the poly-A/RZ module from 172-YTK-P4 using BspHI and EcoRI, blunting the product with T4 polymerase (New England Biolabs). A SmaI site was introduced into GR306 after the SphI site (Figure 3). GR306 was then digested with SmaI, dephosphorylated, and ligated with the blunted poly-A/RZ module, whose correct orientation was confirmed. The resulting plasmid is a multipurpose p1 integration vector in which *mKate2* can be replaced with any target gene. The yeast THI4 coding sequence (*ScTHI4*) was synthesized by GenScript (Piscataway, NJ), and PCR amplified using primers 1 and 2, adding Nsil and SphI sites to the 5’ and 3’ ends, respectively. The amplicon was introduced into pGEM T-Easy by TA cloning (Promega) using the manufacturer’s protocol. *ScTHI4*_pGEM (15 μg) and multi-purpose p1 integration vector (15 μg) were digested with Nsil and SphI, gel-purified, cleaned using a GeneJET Kit (Thermo Scientific), and ligated to give the plasmid p1_ScTHI4 integration vector. In parallel, the multi-purpose p1 integration vector was digested with Nsil and SphI to release *mKate2*, blunted, and self-ligated to generate p1_empty integration vector.

**Figure 3.**
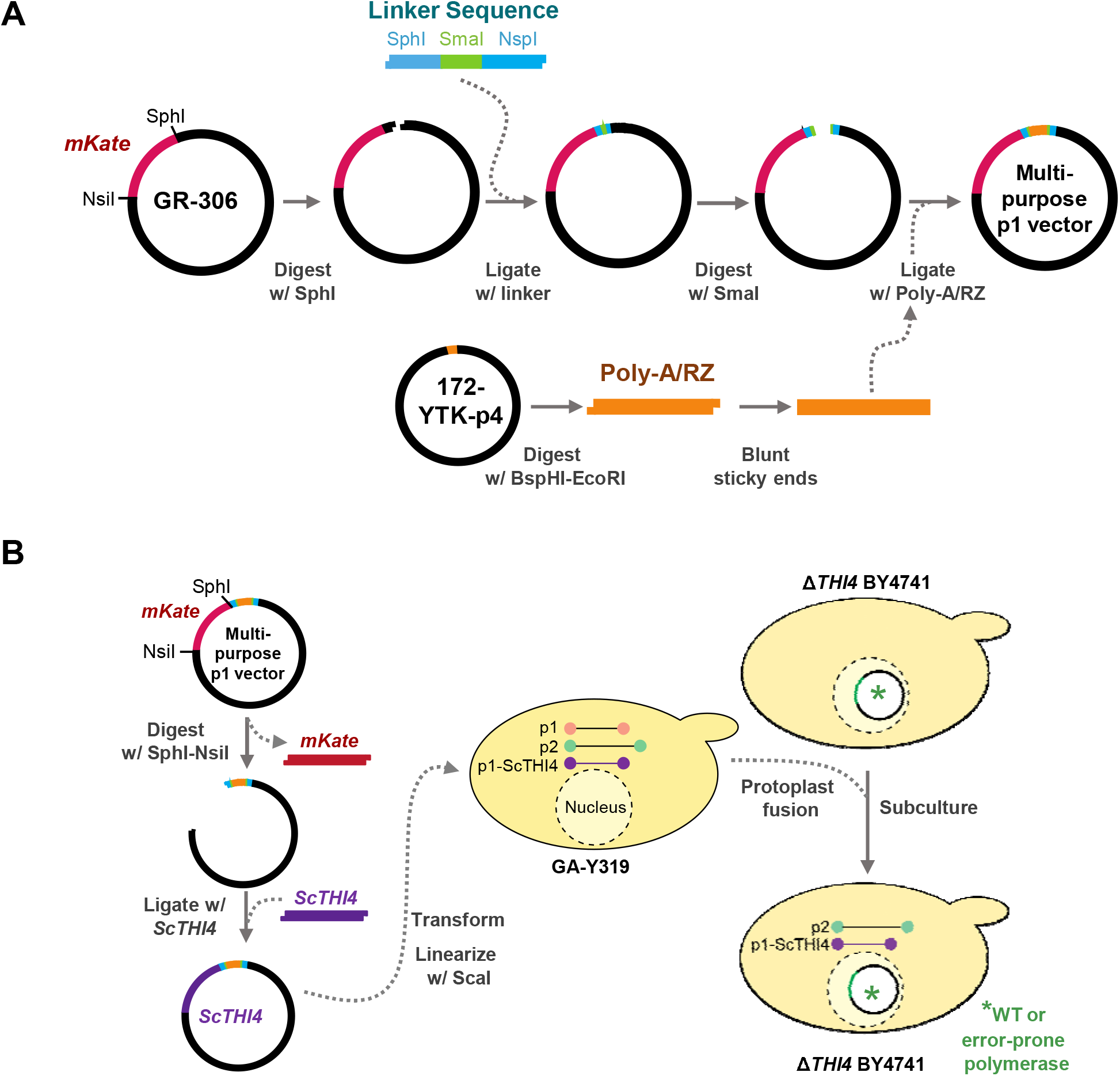
Construction of the multipurpose p1 integration vector for OrthoRep. **(A)** A linker sequence containing a SmaI site was introduced into the SphI site of the GR306 plasmid in such a way as to retain the SphI site. This linker allowed insertion of the poly-A/RZ module from 172-YTK-P4 into GR306. The resulting construct is a multipurpose integration vector from which the resident *mKate* gene can be excised with NsiI and SphI and replaced with a target gene. **(B)** The yeast *THI4* (*ScTHI4)* gene was introduced into the multipurpose integration vector as above; the construct was linearized by ScaI and transformed into yeast strain GA-Y319, and homologous recombination integrated *ScTHI4* into the p1 plasmid. Finally, protoplast fusions were carried out between GA-Y319 (donor strain) and Δ*THI4* BY4741 strains carrying the DNA polymerases TP_DNAP1 wt, TP_DNAP1 611, or TP_DNAP1 633 (receptor strains). Selective medium without leucine, histidine, and tryptophan favored the growth of Δ*THI4* BY4741 cells carrying the full OrthoRep system plus *ScTHI4*.

#### Transformation of the yeast ΔTHI4 strain with wild type and error-prone polymerases

TP-DNAP1 wild type (TP-DNAP1_wt) or error-prone (TP-DNAP1_611 and TP-DNAP1_633) polymerases were cloned into nuclear plasmids containing centromere and autonomous replication sequences (CEN/ARS) and a *HIS3* marker. The resulting plasmids were used to transform the thiamin auxotrophic yeast strain Δ*THI4* BY4741 (*MATa his3*Δ*1 leu2*Δ*0 met15*Δ*0 ura3*Δ*0; THI4*Δ::KanMX) following the PEG-3350/Tris-EDTA/lithium acetate/salmon sperm DNA method with some modifications [40]. The transformants were plated on a synthetic, complete medium (SC; yeast nitrogen base, drop-out mix (USBiological Life Sciences, Cat No. D9540), and 2% (w/v) glucose) minus histidine and supplemented with 100 nM thiamin. Colonies were screened by colony PCR to confirm the presence of the respective TP-DNAP1 sequences (forward primer 3 for all cases and specific reverse primers 4 for TP-DNAP1_wt, 5 for TP-DNAP1_611 and 6 for TP-DNAP1_633).

#### Integration of ScTHI4 into the p1 plasmid

The p1_*ScTHI4* and p1_empty integration vectors (10 μg) were linearized by digestion with ScaI. After inactivation at 80 °C for 20 min, the digest was used to transform strain GA-Y319 (*MAT***a***can1 his3 leu2*Δ*0 ura3*Δ*0 trp1*Δ*0 HIS4 flo1* + p1 + p2) as described [32]. Colonies able to grow in SC medium minus leucine were screened by agarose gel analysis of whole-cell DNA [32] for the presence of both the p1 and p2 plasmids, as well as the successfully recombined p1_*ScTHI4* (Figure S1A) and p1_empty (data not shown).

#### Protoplast fusion of ΔTHI4 BY4741 and GA-Y319 strains

Cultures of recipient (Δ*THI4* BY4741) cells transformed with TP-DNAP1 wild type or error-prone polymerases) and donor (GA-Y319 harboring p1_*ScTHI4*/p2 or p1_empty/p2) strains were grown overnight in YPD medium. Both strains (~0.3 g of cells) were pretreated with a solution of β-mercaptoethanol plus 60 mM EDTA and then incubated with 6 U/mL zymolyase (Zymo Research) in Buffer I to remove the cell wall. The resulting protoplasts were fused in presence of polyethylene glycol-3350 (PEG-3350) and Ca^2+^ (Buffer II) essentially as described previously [41,42]. After incubation for 30 min at 30 °C, the protoplasts were pelleted by centrifugation (10 min, 700 *g*) and gently re-suspended in Buffer III. Finally, cells were embedded in 3% (w/v) agar (dilutions 2:1000 to 2:100) with selective medium maintained at 42 °C (SC minus leucine, histidine and tryptophan supplemented with 0.6 M KCl, 5 g/L ammonium sulfate and 100 nM thiamin), and the plates were incubated at 30 °C. Colonies were recovered from plates and replated on SC medium minus leucine, histidine and tryptophan supplemented with 1 g/L monosodium glutamate, 100 nM thiamin, and 200 μg/mL G418. Individual colonies from plates were then used to inoculate liquid cultures of the same medium to confirm the presence of p1_ScTHI4/p2 and p1_empty/p2 by agarose gel analysis (Figure S1B). The compositions of Buffers I, II, and III are given in Table S2.

#### Functional complementation in yeast

Independent clones of Δ*THI4* BY4741 carrying nuclear plasmids harboring TP-DNAP1_wt, TP-DNAP1_611, or TP-DNAP1_633 and plasmids p1_*ScTHI4* or p1_empty plus wild type p2 were used to inoculate 3 mL of SC minus leucine, histidine, and tryptophan supplemented with 1 g/L monosodium glutamate, 100 nM thiamin, and 200 μg/mL G418 (complementation medium). After 48 h of incubation at 30 °C and 200 rpm, cells were pelleted (3,000 *g*, 5 min) and washed in 5 mL of complementation medium minus thiamin. Aliquots diluted to OD_660nm_ = 0.05 were inoculated in complementation medium plus or minus thiamin and growth was monitored at OD_660nm_.

### EvolvR

#### Bacterial strains and plasmid construction

The *T. ammonificans* THI4 (*TaTHI4*) gene was recoded for *E. coli* and synthesized by GenScript (Piscataway, NJ). The EvolvR plasmid (pEvolvR) encoding nCas9-PolI3M was obtained from Addgene (plasmid 113077). This plasmid has a pBR322 origin of replication and a kanamycin resistance cassette. A tiling approach [43] was used to design a set of guide RNAs targeting different regions of *TaTHI4*, which were cloned into pEvolvR by Golden Gate assembly. Primers are listed in Supplementary Table 1. The pTarget plasmid, p*TaTHI4*, was a pCDF duet vector (Novagen) backbone with a pLlacO1 promoter [44] driving *TaTHI4* and had a streptomycin/spectinomycin resistance marker.

#### Media and culture conditions

The pEvolvR and p*TaTHI4* plasmids were co-transformed into an *E. coli* MG1655 Δ*thiG* strain constructed by recombineering [38]. Three independent clones of each construct were used to inoculate 2 mL of MOPS medium [45] containing 0.2% (w/v) glycerol, 100 nM thiamin, 50 μg/mL kanamycin, and 100 μg/mL spectinomycin. After incubation at 37°C for 18 h, cells were harvested by centrifugation, washed five times with 1 ml thiamin-free MOPS medium and resuspended in 500 μL of the same medium. Liquid cultures of 5 mL of MOPS medium containing 0.2% (w/v) glycerol, 1 mM IPTG, 25 μg/mL kanamycin, 50 μg/mL spectinomycin, and 1 mM cysteine were inoculated with washed cells to an OD_600nm_ of 0.02. Cells were grown at 37°C with shaking at 225 rpm; growth was monitored by measuring OD_600nm_ with a 1.5 cm pathlength cell.

## RESULTS AND DISCUSSION

### OrthoRep

We adapted OrthoRep for high-throughput use by constructing a multi-purpose p1 integration vector based on the GR306 plasmid (Figure 3A), to which we added a convenient directional cloning site for target genes (with Nsil and SphI sites) and a downstream poly-A/RZ module to enhance expression [39]. Because the poly-A/RZ module contains 75 consecutive adenosine residues it is inherently unstable and shortens with repeated propagation. When constructing the multi-purpose integration vector, we verified that ≥72 adenosine residues remained. Note that this vector would need to be reconstructed if further propagation results in failure to reach this threshold.

In continuous directed evolution platforms, target gene function must be coupled to host growth rate. To evaluate the adapted OrthoRep system we therefore tested for complementation of a yeast thiazole auxotroph (Δ*THI4* BY4741) by native *ScTHI4*. We produced Δ*THI4* BY4741 strains harboring the p2 plasmid and combinations of p1_empty or p1_*ScTHI4* with wild type or error-prone (TP_DNAP1_611 or 633) DNA polymerases in the nuclear *CEN6/ARS4* plasmid (Figure 3B). Complementation tests were first run on plates to qualitatively compare the growth of the various strains. The selective medium (minus thiamin, leucine, and histidine) also lacked tryptophan to eliminate any donor GA-Y319 cells (tryptophan auxotroph) left after protoplast fusion (Figure 3B). All strains grew in the presence of 100 nM thiamin and the absence of leucine and histidine (Figure 4A), indicating that all three TP_DNAP1 polymerases were working *in trans* with the recombined p1_*ScThi4*. All three Δ*THI4* BY4741 strains carrying p1_*ScTHI4* grew without thiamin, whereas the empty vector controls did not, as expected. However, the growth rates of the strains carrying p1_*ScTHI4* differed. Cells having the wild type polymerase showed substantial growth after 4.5 days, while cells having TP_DNAP1_611 or 633 did not reach comparable growth until 7 and 8 days, respectively, likely due to the lower copy number of p1_*Sc*THI4 maintained by error-prone TP_DNAP1s, resulting in less expression of THI4. Of the error-prone polymerases, TP_DNAP1_611 gave better growth and so was used for further experiments in liquid medium.

**Figure 4.**
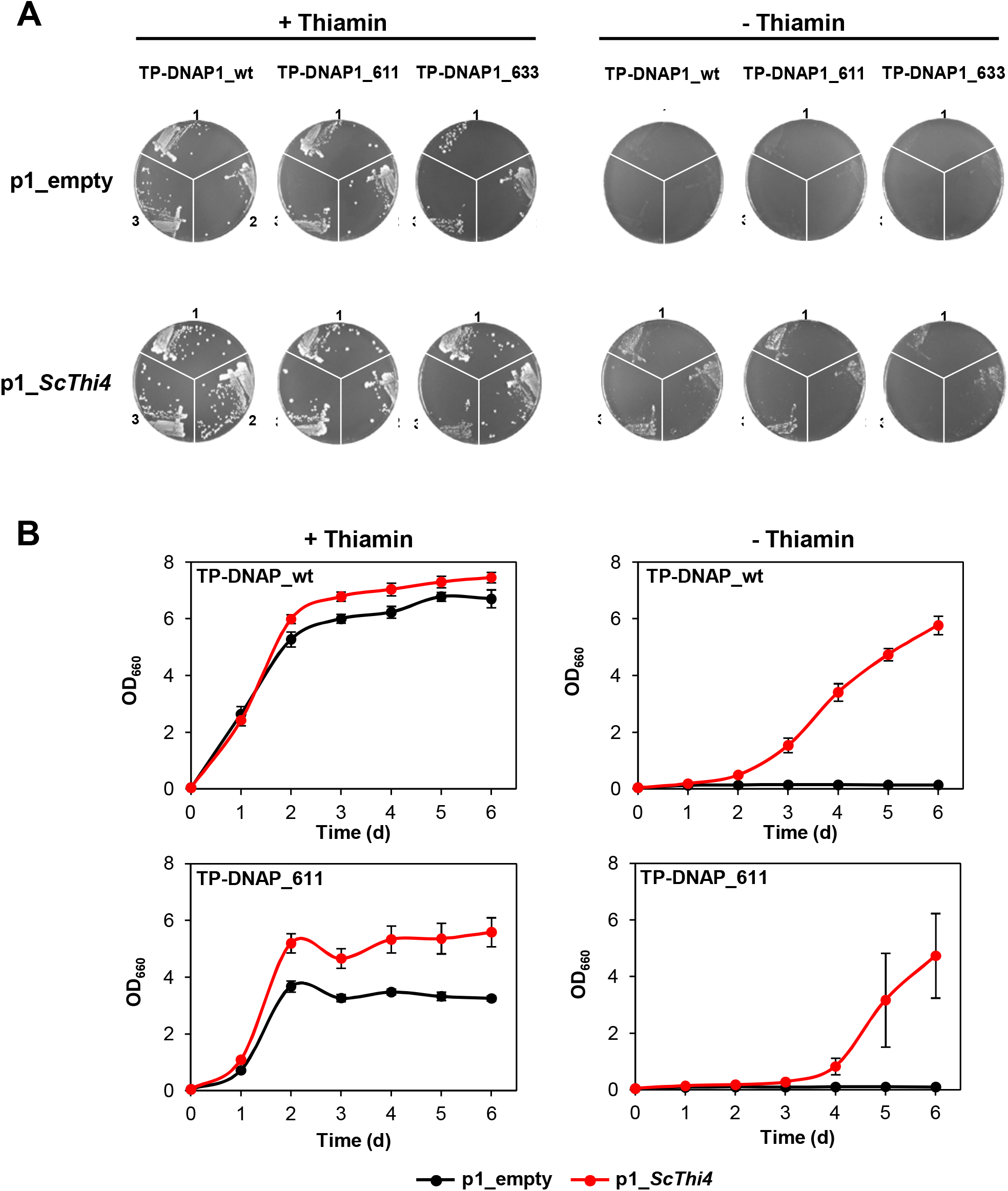
Complementation of a yeast Δ*THI4* strain by the yeast *THI4* gene encoded on the OrthoRep p1 plasmid. **(A)** Three clones of each of the OrthoRep-containing Δ*THI4* strains recovered from protoplast fusions (see Figure 3) were grown on agar plates containing SC minus leucine, histidine, and tryptophan, and supplemented with 2% (w/v) glucose, 1 g/L glutamate and 200 μg/mL G418, in absence or presence of 100 nM thiamin. Clones were collected in 20 μL of the same selective medium (with agar, thiamin, and G418 omitted), from which 3 μL was used to streak fresh plates. Plates containing cells carrying TP_DNAP1 wt, TP_DNAP1 611, and TP_DNAP1 633, were imaged after 4.5, 7, and 8 days at 30 °C, respectively. **(B)** Independent clones from the thiamin-containing plates were used to assess complementation in liquid medium (composition as above, minus agar). Growth of strains containing p1_empty or p1_*ScTHI4*, with or without 100 nM thiamin was measured at OD_660nm_. Values are means ± standard error (SE) of independent replicates (three for TP_DNAP1 611 strains and seven for TP_DNAP1 wt strains). Where no error bars appear, they are smaller than the symbol.

As with solid medium, all strains grew in the presence of thiamin in liquid medium, although the growth of the TP_DNAP1_611 strain was slower than that of the TP_DNAP1_wt strain (Figure 4B). The p1_*ScTHI4* plasmid conferred the ability to grow without thiamin supplementation. However, cells expressing either polymerase showed a growth lag compared to the thiamin-supplemented controls, with the growth of the TP_DNAP1_611 strain delayed more than that of the TP_DNAP1_wt strain and showing more variation among the replicates (Figure 4B). As the copy number of p1_*ScTHI4* is lower in strains with the mutant TP-DNAP1 variants compared to WT, expression of *ScTHI4* is therefore lower and will result in relatively slower growth. Despite the growth lag of the TP_DNAP1_611 strain, the growth period remains short enough (6 days) for continuous directed evolution to be logistically feasible.

### EvolvR

As described in the Introduction, we set out to evolve the non-suicidal TaTHI4 from the bacterium *T. ammonificans* to improve its activity in aerobic, low sulfide conditions similar to those in plant cells. TaTHI4 has a limited ability to complement an *E. coli* thiazole auxotroph under aerobic conditions when supplemental cysteine is given to increase the internal sulfide level [38,46]. Because the *TaTHI4* coding sequence is 0.8 kb, 13 target sgRNAs are needed to cover the whole gene to saturation. We therefore used a tiling approach and designed a set of 13 guide RNAs to target the nickase-polymerase fusion to successive regions of *TaTHI4*. Each pEvolvR construct contained a single guide RNA; we did not concatenate multiple guide RNAs in the same construct because this leads to plasmid instability due to repeated sequence elements. The 13 individual pEvolvR plasmids and the pTarget plasmid harboring TaTHI4 were then co-transformed into the MG1655 Δ*thiG* strain.

To perform directed evolution, we chose the pEvolvR plasmid containing the PolI3M DNA polymerase I variant [27]. First, we checked whether *TaTHI4* expressed from the pTarget plasmid could complement the thiazole auxotrophy of the Δ*thiG* strain in the absence of the pEvolvR plasmid. Cell growth in thiamin-supplemented medium confirmed that *TaTHI4* expression was not toxic (Figure 5A). In medium without thiamin and supplemented with cysteine, the strain carrying the *TaTHI4* pTarget plasmid reached an OD ~2.5-fold higher than the empty vector control, confirming the ability of *TaTHI4* to complement thiazole auxotrophy (Figure 5C). (The initial burst of growth followed by a plateau seen in the empty vector control is attributable to traces of thiazole that contaminate the carbon source or other medium constituents [47]. Active charcoal and ion-exchange pretreatments of the glycerol carbon source were tested in attempts to reduce this unwanted growth but had little or no effect.) In contrast, Δ*thiG* cells carrying the pool of pEvolvR plasmids as well as the *TaTHI4* pTarget plasmid showed no growth in cysteine-supplemented medium, although these cells grew well with thiamin supplementation (Figure 5B, C). This result indicates that pEvolvR expression imposes an insupportable burden on cells cultured in minimal medium, unlike the situation in rich medium [27]. Reducing the copy number of pEvolvR expression level by replacing the high-copy origin (pBR322) with a lower copy variant (p15A), removing non-essential components from the pEvolvR plasmid, or replacing the 0.2% glycerol carbon source with 0.4% or 0.8% glucose did not rectify the problem. Possible future solutions include encoding a single copy of the nickase-polymerase fusion in the *E. coli* genome or modifying the EvolvR system for use in a more robust chassis bacterium such as *Pseudomonas putida* [48].

**Figure 5.**
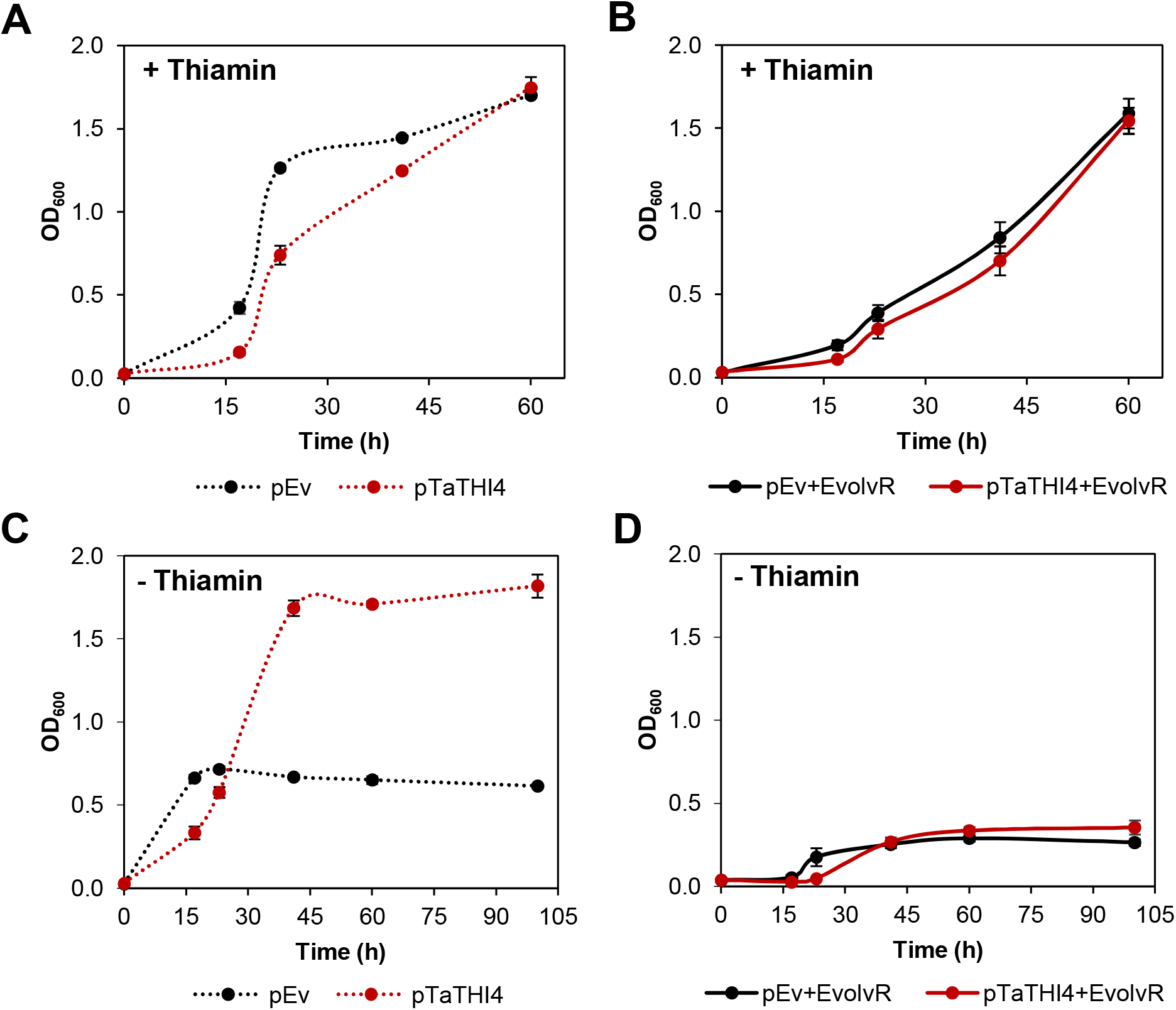
Attempted complementation of an *E. coli* Δ*thiG* strain by *TaTHI4* using components of the EvolvR system. *E. coli* Δ*thiG* cells harboring the pCDFDuet-placO vector alone (pEv) or containing the *TaTHI4* (p*TaTHI4*) gene were cultured in MOPS minimal medium containing 0.2% (w/v) glycerol, 1 mM IPTG, and 1 mM cysteine with 100 nM thiamin **(A)** or without thiamin **(C)**. Δ*thiG* cells harboring both pEvolvR and pEv or p*TaTHI4* were cultured in MOPS minimal medium plus glycerol, cysteine, and IPTG as above, with 100 nM thiamin **(B)** or without thiamin **(D)**. Values are means ± standard error (SE) of three independent replicates. Where no error bars appear, they are smaller than the symbol.

## FUTURE PERSPECTIVES

The above pilot studies indicate that our adapted OrthoRep system is in principle ready for high-throughput evolution of enzymes from plants or other organisms. The EvolvR system, on the other hand, requires modification to overcome the burden of pEvolvR expression in minimal media.

The great potential of OrthoRep and EvolvR lies in their high mutation rates, their scalability, and their compatibility with automation. A major limitation of classical, i.e. discontinuous, directed evolution is that, although a particular suite of mutations may define a fitness peak, the intermediate steps needed to sequentially reach that peak may not survive under strict selection. Automation of OrthoRep can facilitate dynamic selection regimes, as already demonstrated with the real-time/feedback-controlled culture device eVOLVER [29]. By tuning selection stringency, such regimes can help cross fitness valleys [24,25,29]. Moreover, the high mutation rates achievable in OrthoRep and EvolvR favor the accumulation of multiple mutations in a single replication step, which can help skip deleterious intermediate stages as said in the Introduction.

Continuous directed evolution in microbial platform organisms is a TINA (‘There Is No Alternative’) technology as far as plant enzyme improvement is concerned. Implementing directed enzyme evolution in plants is precluded by their large size, long life cycles, and metabolic complexity, which make it infeasible to generate billion-mutant populations and subject them to selection. Taking an enzyme out of the plant, evolving it in a microbe, and reinstalling it in the plant via genome editing is the only practical option. Such a strategy is potentially applicable to five broad areas of plant SynBio, as outlined below and in Figure 6.

**Figure 6.**
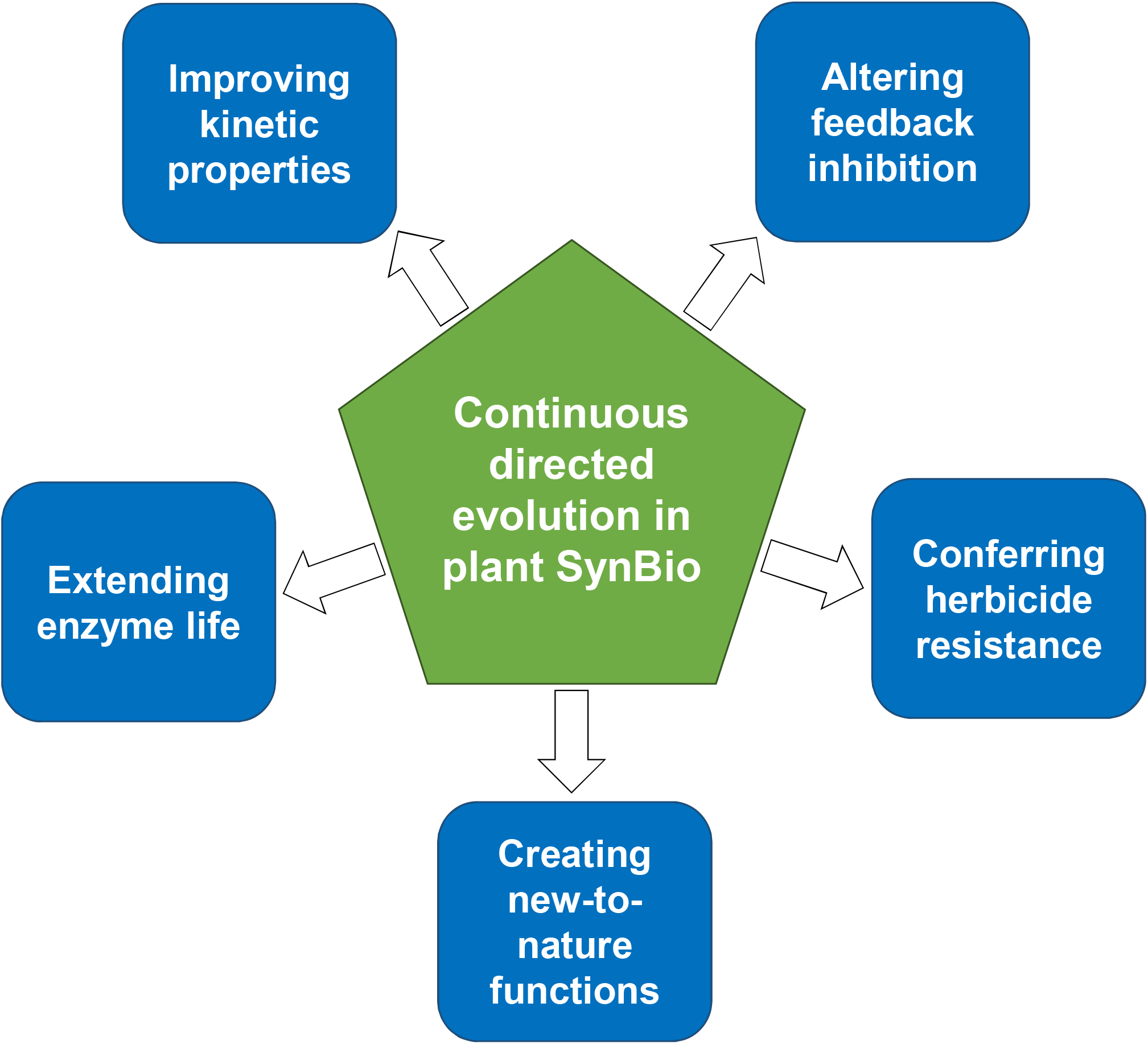
Potential applications of continuous directed evolution in plant SynBio.

### (i) Improving kinetic properties

Directed evolution can outdo rational design by introducing multiple, potentially beneficial changes and going beyond known or predictable key residues [21]. Because it is carried out *in vivo*, continuous directed evolution has the further advantage of improving enzyme function under realistic conditions (e.g. substrate and product levels, macromolecular crowding), which *in vitro* screening protocols may mimic poorly [1].

### (ii) Altering feedback inhibition

Many enzymes at the head of pathways are inhibited by downstream pathway intermediates or end-products and relieving this feedback inhibition is often effective in increasing flux in such pathways [49,50]. Selecting for feedback-insensitive microbial enzymes has a long track record of success [51–53] and can be extended to plant enzymes in microbes engineered to express them [54,55].

### (iii) Conferring herbicide resistance

Many herbicides act by inhibiting essential metabolic enzymes, and mutations that alleviate this inhibition confer resistance [56,57]. Continuous directed evolution has already been used to evolve antibiotic resistance in microbial enzymes [26,27]; this approach could be similarly applied to evolve herbicide-resistant crop enzymes. Using genome editing to put the resistant enzyme back in the crop would avoid using transgenes, which are now banned in various countries [58].

### (iv) Creating new-to-nature functions

Because continuous directed evolution dramatically improves access to the vast protein design landscape, it can create new-to-nature features faster than classic directed evolution [22,59]. Such new features include altered substrate specificity [60] and catalysis of a different type of reaction [8].

### (v) Extending enzyme life

The unnecessarily short working lives of certain plant enzymes wastes resources [3]. Short life is likely due in part to damage to active site residues by chemically reactive substrates or products or by catalytic misfires [61]. A recently introduced metric for enzyme working life – <u>C</u>atalytic-<u>C</u>ycles-till-<u>R</u>eplacement (CCR) = [Metabolic flux rate] / [Enzyme replacement rate] – helps identify enzymes that are candidates for life-lengthening [61]. Continuous directed evolution could potentially be applied to increase enzyme longevity *in vivo*, i.e. to raise CCR, e.g. by driving replacement of damage-prone residues within the active site by less vulnerable alternatives.

## AUTHOR CONTRIBUTIONS

J.D.G. and J.J. performed the experiments, J.J., C.C.R and A.D.H. designed the research. L.T.-R and A.A.J provided experimental support and were supervised by C.R.R. and C.C.L. respectively. A.D.H conceptualized the project and supervised J.D.G., J.J. and J.A.P. J.D.G, J.J., and J.A.P. wrote the initial draft under the supervision of A.D.H. and all authors contributed to review and editing.

