## Supplementary Material for "Potential For Applying Continuous Directed Evolution To Plant Enzymes"

**SUPPLEMENTARY INFORMATION**

**Table S1. Primers used in the present work.**

| **No. of primer** | **Name** | **Sequence 5´- 3´** |
| --- | --- | --- |
| 1 | ScTHI4_Nsil_Fw | CGATATGCATTCTGCTACCTCTACTGCTA |
| 2 | ScTHI4_SphI_Rv | ATCGGCATGCCTAAGCAGCAAAGTGTTTC |
| 3 | TP DNAP1_AREC318_WT_FW | GGTAGACCAAACCATGACTTTGG |
| 4 | TP DNAP1_AREC318_WT_Rv | CCACAATGAGTTCATGATGAT |
| 5 | TP DNAP1_AREC611_ep_Rv | CACAACAATTTATCAAATTCACATTCTAATTCAG |
| 6 | TP DNAP1_AREC633_ep_Rv | GATCAACATGAAGTCTGTATCGTT |
| 7 | TvThi4F1 | TAGCGAGCGAAGTTGTTATTAGCG |
| 8 | TvThi4R1 | AAACCGCTAATAACAACTTCGCTC |
| 9 | TvThi4F2 | TAGCGATGTGGCGATTGTGGGTGG |
| 10 | TvThiR2 | AAACCCACCCACAATCGCCACATC |
| 11 | TvThi4F3 | TAGCGACGTTCGAAGATCGCCACA |
| 12 | TvThi4R3 | AAACTGTGGCGATCTTCGAACGTC |
| 13 | TvThi4F4 | TAGCCGAGATCGTGGTTCAGGAGA |
| 14 | TvThi4R4 | AAACTCTCCTGAACCACGATCTCG |
| 15 | TvThi4F5 | TAGCATTTAAGCCGGGTTACTATC |
| 16 | TvThi4R5 | AAACGATAGTAACCCGGCTTAAAT |
| 17 | TvThi4F6 | TAGCGCGGGTGCGACCGTGTTTAA |
| 18 | TvThi4R6 | AAACTTAAACACGGTCGCACCCGC |
| 19 | TvThi4F7 | TAGCAACGGTCAATACCGTGTGTG |
| 20 | TvThi4R7 | AAACCACACACGGTATTGACCGTT |
| 21 | TvThi4F8 | TAGCTGGTTGGCGAGAAACCGCTG |
| 22 | TvThi4R8 | AAACCAGCGGTTTCTCGCCAACCA |
| 23 | TvThi4F9 | TAGCGAACAGCAAAGAAGTGTTCC |
| 24 | TvThi4R9 | AAACGGAACACTTCTTTGCTGTTC |
| 25 | TvThi4F10 | TAGCCGGTAGCCACCGTATGGGTC |
| 26 | TvThi4R10 | AAACGACCCATACGGTGGCTACCG |
| 27 | TvThi4F11 | TAGCAAAGGTGGCGGAGGAGATTG |
| 28 | TvThi4R11 | AAACCAATCTCCTCCGCCACCTTT |
| 29 | 1944ThermThi4F1F | TAGCGGTTATTACCGCGAAATATG |
| 30 | 1945ThermThi4F1R | AAACCATATTTCGCGGTAATAACC |
| 31 | 1946ThermThi4F2F | TAGCGGTTAGCACCCTGCAGCGTA |
| 32 | 1947ThermThi4R2R | AAACTACGCTGCAGGGTGCTAACC |

**Table S2.** **Compositions of buffers used in the protoplast fusion protocol.** Except for CaCl_2_, all components were diluted in Milli-Q water and autoclaved. A stock of 0.5 M CaCl_2_ was sterilized by filtration (0.22 μm pore diameter membrane) and an appropriate aliquot was added to sterile Buffers II and III.

| Buffer components | Concentration |
| --- | --- |
| **Buffer I**, pH 6.1 |  |
| Citric Acid | 14 mM |
| Na_2_HPO_4_ | 51 mM |
| KCl | 600 mM |
| EDTA dipotassium | 10 mM |
| **Buffer II** |  |
| PEG-3350 | 33% (w/v) |
| KCl | 600 mM |
| CaCl_2_ | 50 mM |
| **Buffer III** |  |
| KCl | 600 mM |
| CaCl_2_ | 50 mM |

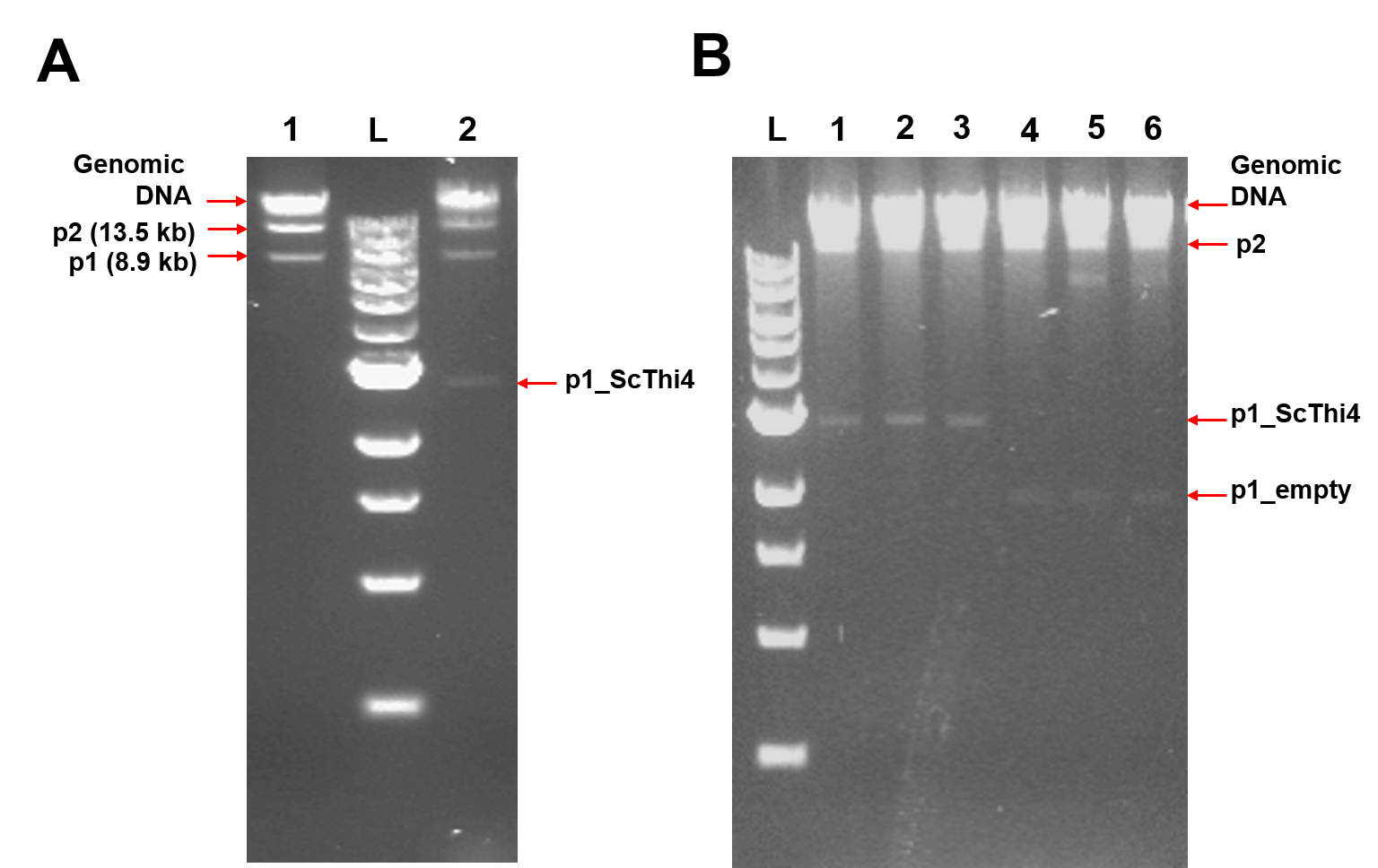

**Figure S1. Gel analysis of DNA extracted from different yeast strains.**

DNA was separated by 1% (w/v) agarose gel. Lanes marked L contained a 1kb DNA ladder.

**(A)** DNA from the GA-Y319 strain contained the wild type p1 (8.9 kb) and p2 (13.5 kb) linear plasmids (Lane 1). After transformation of GA-Y319 cells with a digested p1_*ScTHI4* integration vector, an additional band corresponding to the recombinant p1_*ScTHI4* (3.05 kb) was detected (Lane 2).

**(B)** Screening of Δ*THI4* BY4741 clones after protoplast fusion. DNA from different clones contained the linear plasmids p1_*ScTHI4* (lanes 1-3) or p1_empty (lanes 4-6).
